# Induced pluripotent stem cell-derived human macrophages as an infection model for *Leishmania donovani*

**DOI:** 10.1101/2023.07.31.551225

**Authors:** Lore Baert, Serena Rudy, Mélanie Pellisson, Thierry Doll, Romina Rocchetti, Marcel Kaiser, Pascal Mäser, Matthias Müller

## Abstract

The parasite *Leishmania donovani* is one of the species causing visceral leishmaniasis in humans, a deadly infection claiming up to 40,000 lives each year. The current drugs for leishmaniasis treatment have severe drawbacks and there is an urgent need to find new anti-leishmanial compounds. However, the search for drug candidates is complicated by the intracellular lifestyle of *Leishmania*. Here, we investigate the use of human induced pluripotent stem cell (iPS)-derived macrophages (iMACs) as host cells for *L. donovani*. iMACs obtained through embryoid body differentiation were infected with *L. donovani* promastigotes, and high-content imaging techniques were used to optimise the iMACs seeding density and multiplicity of infection, allowing us to reach infection rates up to 70% five days after infection. IC_50_ values obtained for miltefosine and amphotericin B using the infected iMACs or mouse peritoneal macrophages as host cells were comparable and in agreement with the literature, showing the potential of iMACs as an infection model for drug screening.

**Author Summary:** Yearly, up two million people in poverty-stricken areas contract leishmaniasis, a disease caused by parasites of the genus *Leishmania*. When an infected sandfly takes a blood meal, *Leishmania* parasites enter the host where they are taken up by macrophages. Inside the macrophage, *Leishmania* parasites establish a niche where they can proliferate. Although this infection often leads to disability or death, the drugs currently available are lacking due to toxic side effects, high expenses or difficulties in usage. Drug screening assays that are currently used for compound screening often rely on mouse peritoneal macrophages. We have generated human induced pluripotent stem cell derived macrophages and used these as new host cells for *Leishmania donovani* in the testing of anti-leishmanial compounds. This model has many advantages. For one, it allows us to work with human cells, mimicking the natural infection more closely than possible with murine cells. Secondly, it allows to obtain bigger batches of uniform cells for screening campaigns. Finally, this approach aligns with the principle of 3R, replacing the use of animals for cultivation of *Leishmania* and drug screening purposes.

## Introduction

Up to one million people in nearly 100 endemic countries contract leishmaniasis every year, rendering this often fatal or debilitating infection one of the most important neglected tropical diseases (1). Leishmaniasis is caused by protozoan parasites of the genus *Leishmania*, which are transmitted to a mammalian host when an infected phlebotomine sandfly takes a blood meal (2). Upon entry into the host, the parasites are phagocytosed, mostly by macrophages. Strikingly, *Leishmania* parasites not only survive but even thrive in the hostile environment of the macrophages, where they establish a niche in the parasitophorous vacuole (3,4). *Leishmania* is characterised by two distinct life cycle stages. Inside the sandfly, *Leishmania* exists as its promastigote form, which is recognisable by a single, long flagellum protruding from the anterior end of the cell body. Once inside the macrophages, the acidic environment and higher temperature trigger the differentiation of the promastigotes to amastigotes, which are smaller and only have a rudimental flagellum (4,5).

More than 20 known *Leishmania* species have the ability to infect humans, causing different forms of leishmaniasis (6). The most severe form is visceral leishmaniasis (VL), an infection of the internal organs, in particular the liver and the spleen. VL is caused mainly by *L. donovani* and *L. infantum* and is almost always fatal if left untreated (7). While there are effective drugs on the market, they all have severe drawbacks regarding either high cost, the impracticability of use, adverse side effects, or rising drug resistance (8). Thus, there is an urgent need for new and better treatments options.

The intracellular lifestyle of *Leishmania* complicates the search for new active molecules. Compounds are most relevantly tested against the intracellular amastigotes (9). Macrophages obtained from various sources have been used to study host-parasite interaction and anti-leishmanial drug efficacy. These include bone marrow-derived murine macrophages (10), peritoneal murine macrophages (PMMs) (11,12), human peripheral blood monocyte-derived macrophages (13), and immortalised monocyte-like cell lines such as THP-1 cells (13,14). Each of these has some disadvantages. The primary macrophages are incompatible with high-throughput screening as their isolation is time-consuming, the yields are low, and there is a large variation between batches(15). On the other hand, the malignant origin of the monocyte-like cell lines such as THP-1, makes them genetically unstable and therefore unreliable for drug screening and investigating host-parasite interaction (16). A promising alternative to these macrophage populations are iPS-derived macrophages (15,17–19).

In 2012, Shinya Yamanaka and John B. Gurdon received the Nobel Prize for their discovery that somatic cells can be dedifferentiated into a pluripotent state by overexpression of the transcription factors Oct4, Sox2, Klf4 and c-Myc (20). Since then, induced pluripotent stem cells (iPSCs) have been differentiated into dozens of different cell types, including hepatocytes (21,22), neurons (23,24), endothelial cells (25) and macrophages (18,19,26). iPS-derived macrophages (iMACs) highly resemble their counterparts isolated from human donors, making them a favoured system compared to transformed cell lines like THP1, which often harbour multiple uncharacterised genetic mutations (27). Working with human iMACs versus primary murine cells is not only more practical but also more ethical as it reduces the use of laboratory animals. In addition, the use of human iMACs is more relevant since mice have a different immune profile than humans. For example, even in susceptible strains such as the BALB/c mice, visceral leishmaniasis is not fatal, which is in sharp contrast to infection in humans (28,29). Finally, it is relatively easy and quick to generate large quantities of genetically identical iMACs. All these advantages make iMACs an attractive model for drug screening.

In this study, we made use of human iMACs to establish a new infection model for *L. donovani*. After optimising the culture condition for infectibility with late-stage promastigotes the two reference drugs, miltefosine and amphotericin B, were tested for their anti-leishmanial activity in iMACs and compared to the standard model using PMMs.

## Materials & Methods

### Media

*Leishmania culture medium:* Equal parts SM and SDM-79 medium (30,31) (pH 7.4) supplemented with 10% heat-inactivated FCS and 2 ug/mL hemin. *mTeSR Plus medium:* mTeSR Plus Basal Medium (STEMCELL Technologies) supplemented with mTeSR Plus 5x Supplement and 1% penicillin/streptomycin (Gibco). *Complete RPMI medium:* RPMI 1640 Glutamax (Gibco) supplemented with 10% heat-inactivated FCS (Gibco), 1% sodium pyruvate (Gibco), 1% penicillin/streptomycin, 25 mM HEPES (Gibco), 0.055 mM β-mercaptoethanol (Gibco). *MDM medium:* DMEM (Gibco) supplemented with 10% FBS, 1% Glutamax (Gibco), 0.055 mM β-mercaptoethanol, 1x MEM Non-Essential Amino Acids (Gibco) and 1% penicillin/streptomycin. *X-Vivo 15 medium:* X-Vivo 15 (Lonza) supplemented with 1% Glutamax, 0.055 mM β-mercaptoethanol and 1% penicillin/streptomycin.

### *L. donovani* culturing

*Leishmania donovani* (MHOM/ET/67/HU3) promastigotes were maintained at 27°C, 5% CO_2_, in *Leishmania* culture medium. Cultures were passaged three times a week (1:100). To obtain metacyclic promastigotes, the cultures were incubated for 5 days until reaching stationary growth phase.

### iPS cell culturing

Female human WT29 iPS cells were derived from cell line AG092429 obtained from the NIA Aging Cell Repository at the Coriell Institute for Medical Research. The male iPS line WT94 was generated from GM23394 also obtained from Coriell. Stem cells were plated on a LN511-coated surface (BioLamina) and amplified in mTeSR Plus medium. During the first 24 h after seeding, the medium was supplemented with 10 µM ROCK Inhibitor (Y-27632). The medium was changed every 24h and the cells were passaged every 3 days (1/30). For passaging, the cells were washed with PBS and detached by incubation with TrypLE (Gibco) for 5 min at 37°C.

### iMAC differentiation

On day 1, iPS cells were resuspended in mTeSR Plus medium supplemented with 10 µM ROCK Inhibitor at a concentration of 60,000 cells/ml. The iPS cells were distributed to BIOFLOAT 96 well plates (FaCellitate) at 100 µL/well. On day 2, 100 µL mTeSR Plus medium supplemented with 100 ng/mL BMP4, 40 ng/mL hSCF and 100 ng/mL VEGF was added to each well. On day 3, 100 µL of medium was replaced with fresh mTeSR Plus medium supplemented with 50 ng/mL BMP4, 20 ng/mL hSCF and 50 ng/mL VEGF. On day 5, embryoid bodies (EBs) measuring 400 to 700 nm in diameter were apparent. These were resuspended in MDM medium supplemented with 100 ng/mL hm-CSF and 25 ng/mL IL3. The EBs were then plated in a T75 flask coated with 0.1% gelatine in PBS. On day 8, the medium was replaced with X-Vivo 15 medium supplemented with 100 ng/mL hm-CSF and 25 ng/mL IL3. Every 3 to 4 days, two-thirds of the medium were replaced with fresh medium. About 3 weeks later, iMACs were harvested from the supernatant for about three months. Incubation of the cells was done at 37°C, 5% CO_2_.

### iMAC polarisation

After harvesting, iMACs were centrifuged for 5 min at 200g and resuspended in Complete RPMI medium supplemented with 40 ng/mL M-CSF and seeded in 96 well plates. For polarisation to M1 or M2 states, 50 ng/mL IFNγ and 50 ng/mL Dexamethasone or IL-4 was added to the medium, respectively. The polarisation was done for 48h or 5 days. Growth factors were maintained in the medium during the entire assay.

### iMAC characterisation using flow cytometry

iMACs were either directly harvested from the EB culture or detached by incubation with TrypLE for 5 min at 37°C following polarisation. iMACs were then washed with PBS plus 2% FBS and incubated for 15 min at RT with 50 ul/mL Human TruStain FcX Blocking solution (BioLegend) in PBS + 2% FBS. Next, iMACs were stained for 1h on ice with mouse anti-human antibodies CD11b-FITC, CD11c-PE, CD14- PE/ AF488, CD16-FITC, CD45-PE/ FITC, CD68-FITC, CD80-FITC, CD86-AF488, CD163-FITC, CD206-AF488 and the corresponding IgG1κ or IgG2b isotype controls (Invitrogen). Staining was followed by two washing steps in PBS + 2% FBS, and fixation of the iMACs for 15 min at RT in BD Cytofix buffer (BD Biosciences). Finally, the samples were washed twice before being resuspended in PBS plus 2% FBS. The Attune NxT Acoustic Focusing Cytometer was used for measuring the samples (2.0*10^4^ events/sample). The data were analysed using FlowJo software.

### Murine cell culturing and seeding

Two days prior to harvesting of PMMs, CD1 mice were inoculated with 0.5 mL starch solution (Fluka/ Sigma 85645). To harvest the cells, mice were euthanised and 10 mL of RPMI medium was injected into the peritoneal cavity using a 25G needle. Next PMMs were extracted from the peritoneal cavity using a 22G needle. The PMMs were centrifuged 10 min at 200g and resuspended in RPMI medium containing 1% Mäsermix (32) complemented with 15% RPMI medium containing growth factors obtained through 7 day cultivation of the LadMac cell line. Subsequently the PMMs were seeded in T75 flasks and incubated 37°C, 5% CO2. After 3 days, the PMMs were washed with EBSS and trypsinised for 20s using 0.05% trypsin. Using a cell scraper, the PMMs were fully detached and collected in complete RPMI medium before seeding in flat bottom cell culture microplates (uClear, Greiner Bio-One, Cell Star) for infection.

### Infection with *L. donovani* and drug treatment

On day 1, iMACs or PMMs were seeded at various densities in flat bottom cell culture microplates (uClear, Greiner Bio-One, Cell Star) in Complete RPMI medium plus 40 ng/mL CSF. The outer wells of the plates were not used and filled with PBS. On day 3, the medium was aspirated. Late-stage promastigotes were centrifuged (15 min at 800 g), resuspended in fresh Complete RPMI medium plus 40 ng/mL CSF, and added to the macrophages at different MOIs. On day 4, extracellular parasites were washed away twice with medium before fresh Complete RPMI medium + 40 ng/mL CSF was added to the macrophages. When drugs were used during the assay, they were added at this point. Either miltefosine (M5571, Sigma-Aldrich) resuspended in H_2_O or amphotericin B (A2411, Sigma-Aldrich) resuspended in DMSO were added to the medium at various concentrations (final concentration of DMSO <0.05%). On day 8, the macrophages were fixed and stained. Macrophages were always incubated at 37°C, 5% CO_2_.

### Fixation and staining

Macrophages were fixed with 4% formaldehyde for 20 min at RT in the dark (Sigma). After fixation, the cells were washed three times with PBS.

### DAPI and AF488-phalloidin stain

Macrophages were permeabilised with 0.1% Triton X-100 diluted in PBS for 90s. Afterwards, the cells were washed once with PBS. Next, the macrophages were stained with 100 µL of 5 µg/mL DAPI in PBS for 10 min at RT in the dark. Another washing step was performed with PBS before staining with 0.033 µM AF488 phalloidin (BioStatus) for 30 min at RT in the dark. After staining, the macrophages were washed three times with PBS. The wells were filled up with PBS and the plates were stored at 4°C.

### DRAQ5 stain

Macrophages were stained with 50 µL of 5.0 uM DRAQ5 (BioStatus) in PBS for 20 min at RT in the dark. After staining, the macrophages were washed three times with PBS. The wells were filled up with PBS and the plates were stored at 4°C.

### LAMP-1 stain

iMACs were permeabilised with methanol for 20 min at -20°C. Afterwards, cells were incubated for 1h in 0.1% Triton X100 and 2% BSA (Sigma Aldrich) diluted in PBS. This was followed by overnight incubation at 4°C with 1:200 anti-LAMP-1 (D401sm #15665, Life Technologies). The next day, the iMACs were washed 3 x 5 min with PBS and incubated for 1h with 1:1000 goat anti-mouse IgG-AF488 (A11001, Invitrogen). Afterwards the cells were again washed 3 x 5 min with PBS. The wells were filled up with PBS and the plates were stored at 4 °C.

### Imaging and analysis

Infection assays were either imaged using the ImageXpress Micro XLS High Content Imager (Molecular Devices) or the CV7000 High Content Imager (Yokogawa), depending on availability. Between 9 to 16 images per well were taken at 20x objective lens ensuring at least 13% surface coverage. MetaXpress (Molecular Devices) or CellPathfinder (Yokogawa) software was used for automatic analysis of the images including counting the number of macrophages, the percentage of infected macrophages, and the number of intracellular *Leishmania*. iMACs stained with LAMP-1 antibody were imaged using confocal microscopy. Images were taken at 40x objective lens using the LSM900 Airyscan (ZEISS) and analysed using the ZEN 3.2 software (ZEISS). For the calculation of the percentage of infected macrophages and the number of intracellular parasites per infected macrophage, the counts were corrected by subtracting the false positive signal calculated from an uninfected negative control sample. When performing drug assays, an untreated positive control sample was included and the percentage of infected macrophages and number of intracellular parasites per macrophage were corrected with the negative control and normalised by dividing the values with the corrected positive control value and multiplying by 100.

### Statistics

Data analysis was done using GraphPad Prism 9.5.1 software. All data are shown as mean ± SD calculated from a minimum of three independent experiments. IC_50_ values of compounds were calculated using non-linear regression to a sigmoidal dose-response curve. Statistical analysis was done using ANOVA tests and t-tests. P-values < 0.05 were considered statistically significant.

## Results

### Generation of iPS-derived macrophages

iMACs were generated according to the published protocol of van Wilgenburg *et al* (18). Two to three weeks after inducing the formation of embryoid bodies (EBs) from human iPS cells (line WT29), the EBs started to produce iMACs of the M0-like subtype in the supernatant expressing CD11b, CD14, CD16, CD45, CD68 and CD163 (Fig. 1A and Fig. S1). To test the plasticity of iMACs they were polarised with IFNγ or dexamethasone into either the inflammatory M1 or anti-inflammatory M2c subtypes, respectively. On top, 2 or 5 days treatment was tested to evaluate whether a shorter polarisation time suffices. Polarisation success was assessed by measuring the expression of the subtype-specific macrophage markers CD80 (M1) and CD163 (M2) (Fig. 1B). As expected, CD14, a marker for cells of the myeloid lineage, was highly expressed over all conditions. The M1 marker CD80 showed a high expression level in M1 iMACs but was undetectable in other conditions. Expression of the M2 marker CD163 was present in all subtypes but was obviously higher in M2c iMACs. There was no striking difference in expression patterns for the M1 and M2 specific CD80 and CD163 markers between iMACs that had been polarised for 2 vs. 5 days suggesting a shorter polarisation time suffices. This demonstrates that iMACs can response to inflammatory stimuli like primary macrophages (15).

**Fig. 1:**
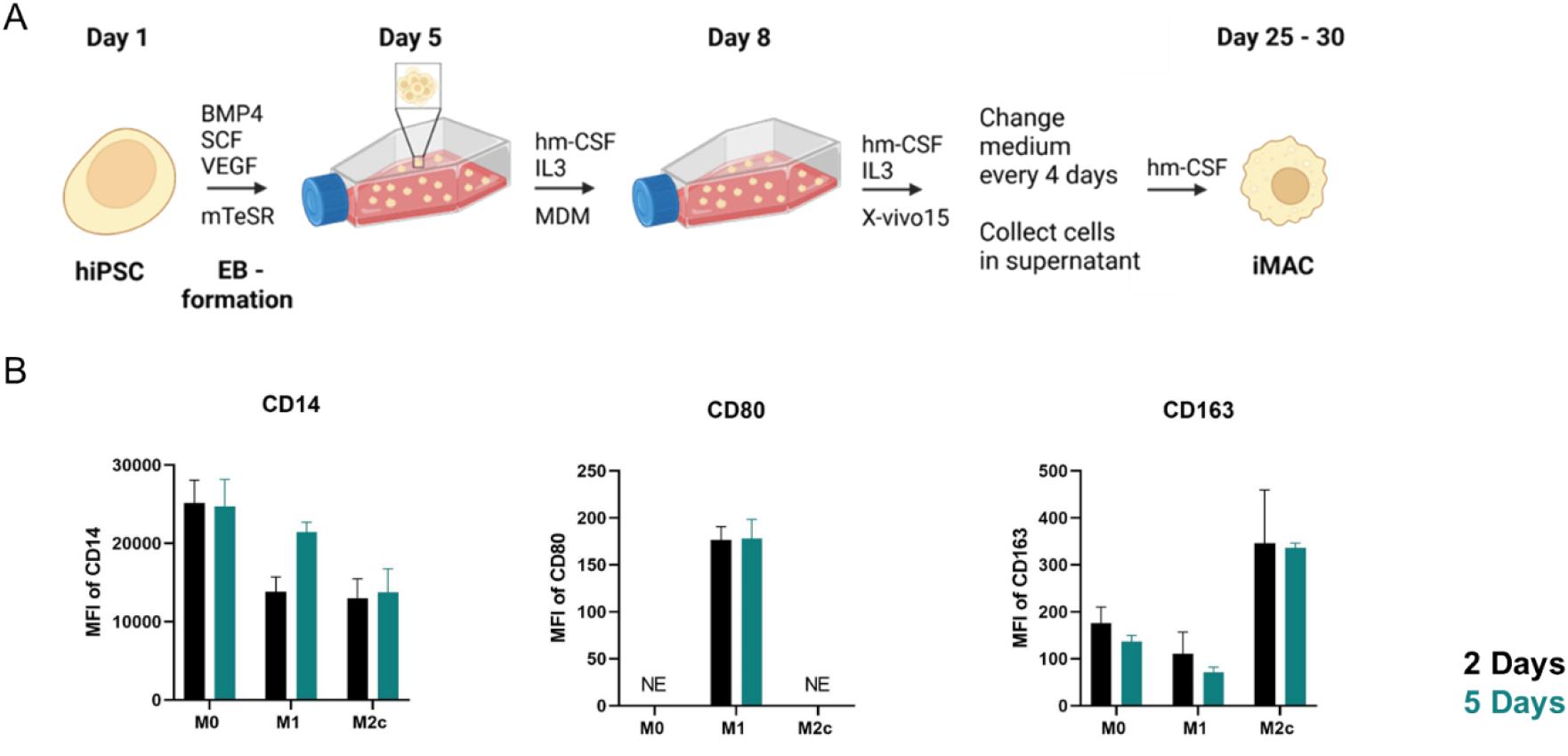
Differentiation of iPS cells to iMACs followed by polarisation. A) Schematic overview of the differentiation process of iPS cell to macrophage (iMAC) via the formation of embryoid bodies (EBs) B) iMACs were polarised to the M0, M1 or M2c subtypes over either 2 (black) or 5 days (green). Expression of the cell surface receptors CD14 (left), CD80 (middle) and CD163 (right) was measured by flow cytometry. The data is shown as the MFI (geometric mean) ± SD, normalised by the isotype control. N = 3, NE = Not Expressed.

### Determination of optimal experimental conditions for infection of iMACs with *L. donovani*

In order to obtain infection rates high enough to provide a window for drug screening in 96 well plates, we determined the optimal parameters for an *L. donovani* infection model using iMACs as host cells. For this purpose, *L. donovani* promastigotes were grown until the late stationary growth phase in order to obtain a higher proportion of infective metacyclic parasites, which are recognisable by their long and slender appearance (Fig. 2A). Late stationary phase was reached after 5 days of incubation starting with a population of 1.0E+05 cells/ mL (Fig. 2B). For infection, iMACs were seeded at densities of 10,000 to 60,000 iMACs per well and infected with *L. donovani*. To test the infection methodology, an MOI of 20 was taken, based on the current in-house infection assay done with PMMs. When the seeding density of iMACs was higher than 20,000 cells per well, there was a significant formation of cell clumps (Fig. 2C). This severely interfered with the automated image analysis used for counting the number of iMACs and intracellular parasites. Indeed, with increasing seeding densities, the iMAC count reached a plateau at about 700 iMACs per image when more than 20,000 iMACs were seeded (Fig. 2D). The same trend was seen for both infected and uninfected iMACs. Based on these findings, we concluded that the seeding density should not surpass 20,000 iMACs per well, which resulted in an average cell count of 435 ± 80 iMACs per image.

**Fig. 2:**
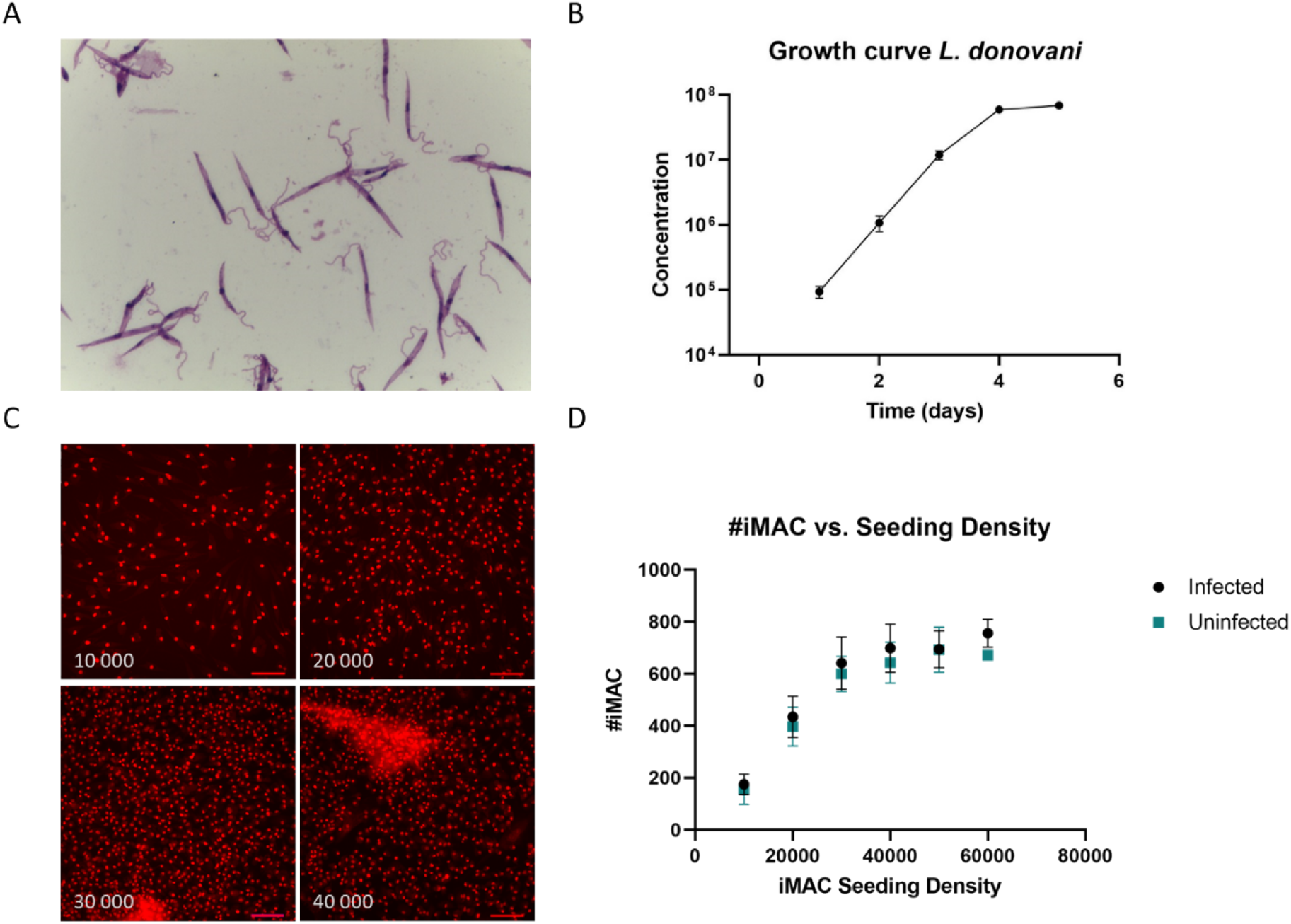
Growth characteristics of Leishmania donovani and optimal iMAC seeding density. A) Giemsa staining of late stage stationary promastigotes taken after 5 days in culture B) Growth curve of *L. donovani*, with a starting concentration of 10^5^ parasites/mL, N=3 C) Representative images of uninfected iMACs seeded at densities of 10,000 to 40,000 cells/well, stained with DRAQ5. Scale bar = 100 um. D) Average number of iMACs/image in function of the initial seeding density for infected (black) and uninfected (green) iMACs. N=4

Since an MOI of 20 resulted in low infection rates, we tested the infection of iMACs with MOIs between 50 and 300 and assessed the infection rate 5 days later. We found that with an increasing MOI, more iMACs were lost (Fig. 3A, B). It is currently unclear whether this effect is due to the rupture of infected macrophages or simply to their detachment and loss during media change. At all MOIs tested, we observed a high infection rate, which plateaued at 75.0 ± 10.2% at an MOI of 200. A similar pattern was found for the number of intracellular parasites, with an average of 5.6 ± 2.0 parasites per infected iMAC, plateauing at an MOI of 250 (Fig. 3C). However, a considerable number of cells harboured up to 50 parasites when infected at higher MOIs (Fig. 3D). Based on these results, we decided to use an MOI of 100. With this MOI, 65.9 ± 13.9% of iMACs are infected and 82.5% of them have up to 10 *Leishmania* parasites per infected cell. To test the robustness of the infection model, iMACs were generated from an additional male iPS line (WT94) and infected at an MOI of 100. No significant difference was found between infection rates of WT29 and WT94 at 64.8 ± 10.4% and 69.7 ± 11.7% respectively (Fig. 3E). The average number of *Leishmania* per iMAC was also not significantly different between both cell types. Taken together, these results suggest that iMACs are a suitable model for *L. donovani* infection.

**Fig. 3:**
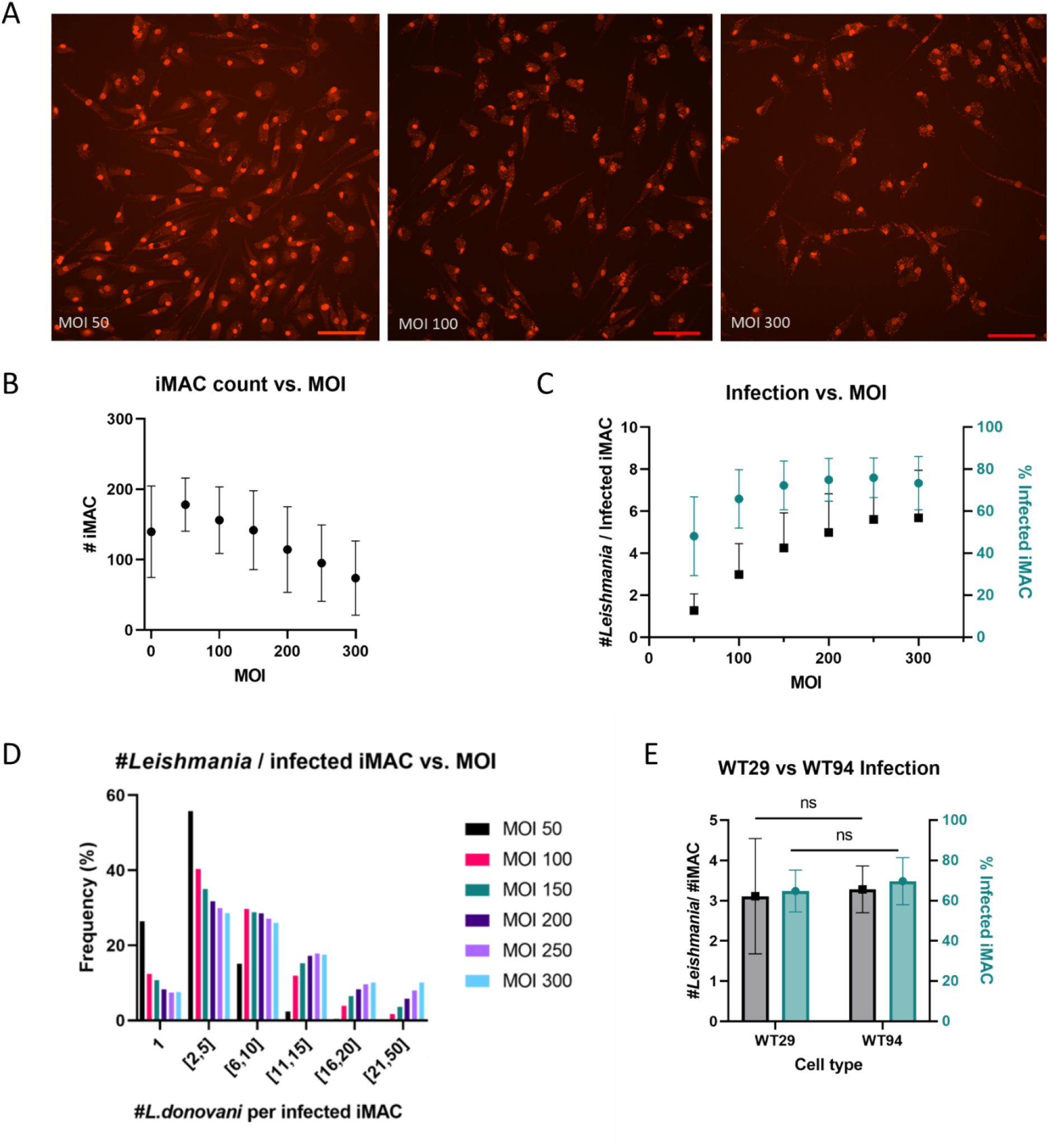
iMAC count and infected iMACs (%) vs. parasite MOI. A) Representative images of iMACs infected with *L. donovani* at a MOI of 50, 100 and 300, stained with DRAQ5. Scale bar = 100 um. B) Average number of iMACs/ image in function of the MOI. C) iMAC infection in function of MOI, presented as the number of intracellular parasites per infected iMAC (black) and the percentage of infected iMACs (green). The infection rate was normalised by subtracting the value of the negative control. N=3 D) Histogram showing the frequency distribution of number of intracellular *L. donovani* per infected iMAC in function of the MOI. E) Infection of WT29 and WT94 iMAC, presented as the number of intracellular parasites per infected iMAC (black) and the percentage of infected iMACs (green). N = 3. *P<0.05 was considered significant using unpaired t-test.

### Influence of iMAC age and polarisation on infection rate

Once an iMAC-producing culture was established, iMACs could be harvested at each medium change over a period of 3 to 4 months. Afterwards the production of the iMACs was markedly reduced and was discontinued. To see if early-time versus late-time iMACs are comparable, the surface markers and infection rate were tested at two different time points. The expression of the CD markers (CD11c, CD14, CD45, CD68, CD86, CD163 and CD206) were comparable between iMACs harvested from young or old cultures, which is consistent with other publications (15) (Fig S2a). To check the iMAC infectivity, iMACs harvested from young (<1.5 months) and old (>2.5 months) EB cultures were infected *with L. donovani* at an MOI of 100. Analyses 5 days after infection could not detect any significant differences between the two conditions (Fig. S2b). The age of the culture therefore appears to have little effect on infection rate.

As M1 macrophages are inflammatory while M2 macrophages work anti-inflammatory and help with tissue repair, it is possible that the macrophage subtype influences infectibility of the iMACs with *L. donovani*. We polarized M0 macrophages to either the M1 or M2c subtype and infected them with *L. donovani* at a MOI of 100. There is no observable difference between the three subtypes 5 days after infection (Fig. S3). Both the number of parasites per infected iMAC and the percentage of infected iMACs were comparable over all conditions (Fig. S3).

### Localisation of Leishmania in the iMACs

Based on these results, we seeded M0 iMACs with a density of 10,000 iMACs per well and infected with a MOI of 100. Staining was done with a DAPI-phalloidin double stain (Fig. 4A) or a DRAQ5 stain (Fig. 4B). Both stains allowed a clear visualisation of the iMACs and the intracellular *L. donovani* parasites. Z-stacking using confocal microscopy clearly showed how the parasites resided within the iMACs and were not e.g. sticking on the outside of the cells (Fig. 4C). To determine intracellular localisation of the parasites, we stained the infected iMACs with Lysosomal-associated membrane protein 1 (LAMP-1) antibodies, a lysosomal marker. Indeed, we observed a localisation of *L. donovani* inside vacuoles that recruit LAMP-1, pointing towards an intralysosomal localisation of the parasites (Fig. 4D).

**Fig. 4:**
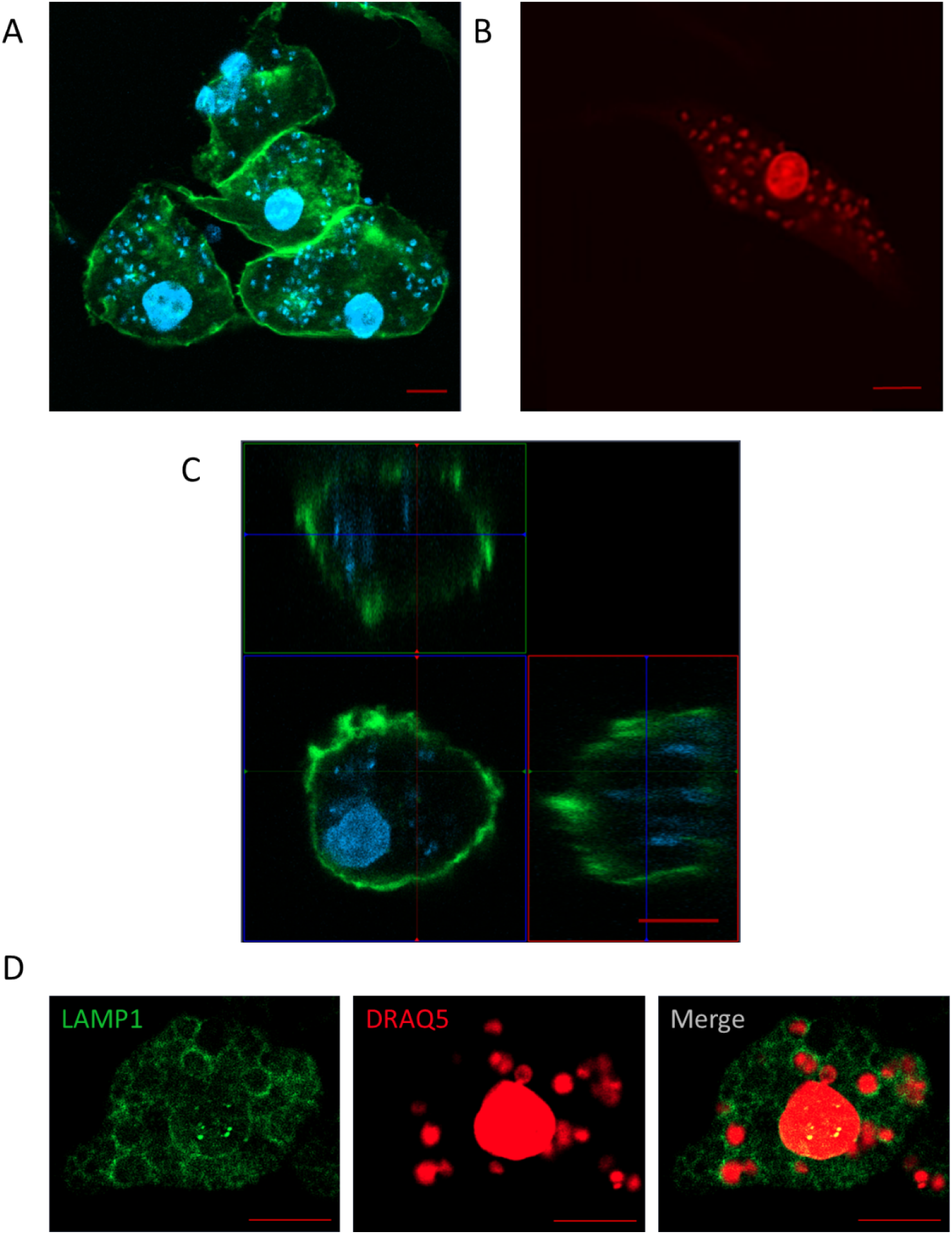
Characterization of iMACs infected with L. donovani. Infected iMACs were stained with A) DAPI (blue) and AF488 phalloidin (green) or B) DRAQ5 (red) C) Intersection of infected iMAC infected with DAPI (blue) and AF488 phalloidin (green) D) LAMP1 staining of infected iMAC. Scale bar = 10 um

### Testing the iPS-derived system for drug susceptibility

Finally, we tested the reference drugs miltefosine and amphotericin B against infected iMACs and calculated the 50% inhibitory concentration (IC_50_) values based on the decrease of infected iMACs or decrease of number of intracellular parasites per iMAC. As control, we also tested these reference drugs against infected PMMs. Cells were treated for 4 days with miltefosine (0.16 to 40 uM) or amphotericin B (0.82 to 200 nM) 24h after initial infection. As high concentrations of miltefosine are toxic to the macrophages, we left out the highest concentration (40 uM) out of the analysis. There was a clear decrease in parasitemia over all conditions (Fig. 5). The IC_50_ values calculated based on the number of intracellular parasites were consistently lower than those based on the percentage of infected iMACs (Table 1). Looking at the former parameter, the IC_50_ for miltefosine using WT29- derived iMACs as host cells was 3.85 ± 1.90 µM, which was comparable to what had been described in literature and to the IC_50_ calculated for the PMM-model (2.80 ± 0.99 µM) (33). For amphotericin B, we calculated an IC_50_ of 62.1 ± 39.0 nM using WT29-derived iMACs. The IC_50_ for the PMM model was lower with 12.8± 4.76 nM. Both values were in the range of what had been reported before (34). Repeating the same experiment with iMACs generated by iPS line WT94 leads to very comparable results. The IC_50_ value calculated from the value for infected cells was 3.52 ± 2.28 µM for miltefosine and 58.0 ± 28.2 nM for amphotericin B (Table 1). This high correlation between WT29 and WT94 indicates that iPS-derived macrophages could be a valuable tool in the search for new drugs.

**Fig. 5:**
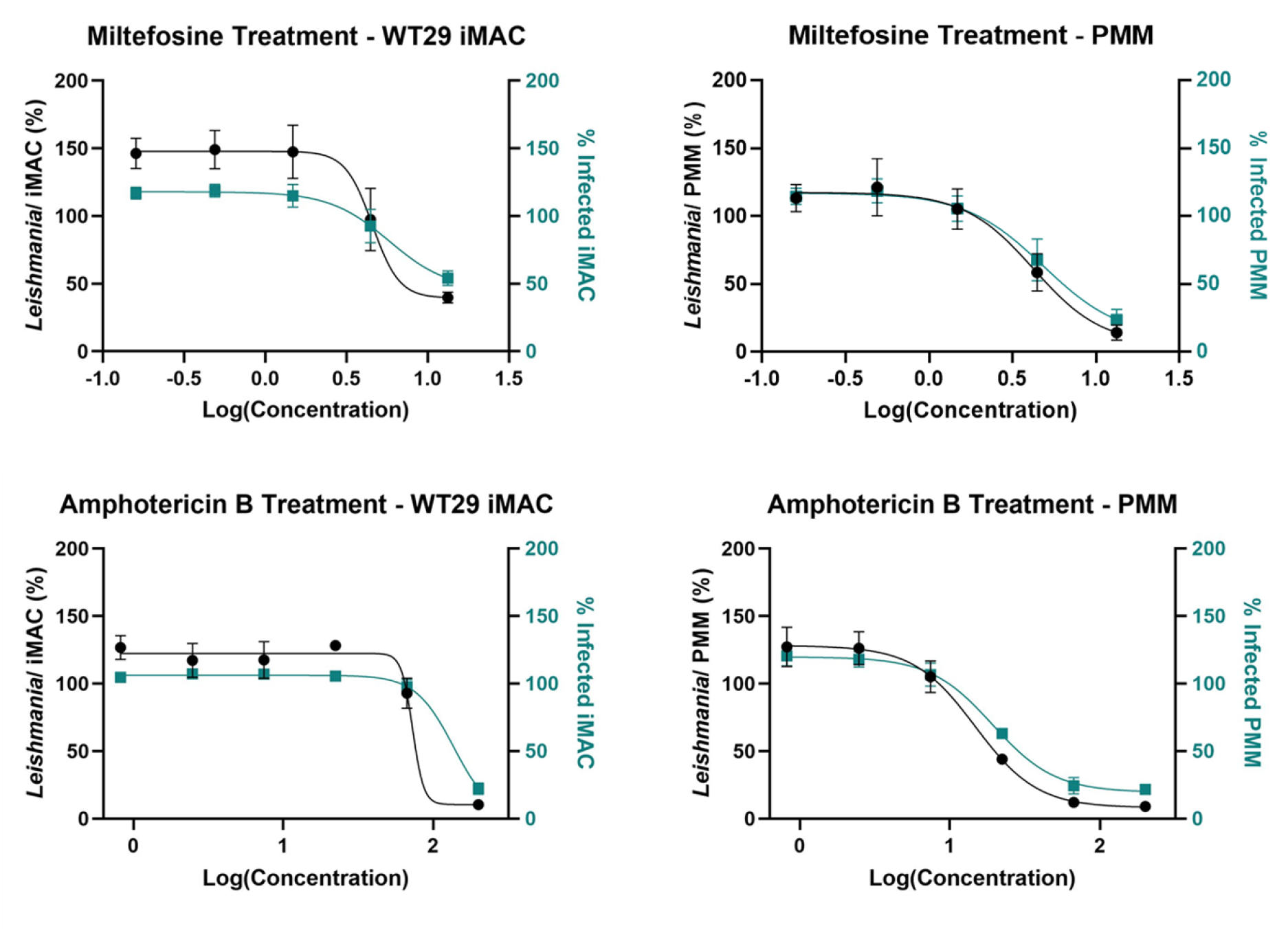
Drug response for miltefosine and amphotericin B. Infected WT29 iMACs and PMMs were treated with miltefosine or amphotericin B. The number of intracellular *L. donovani* parasites per cell for each concentration is shown on the left y-axis (black). The average number of infected cells is shown on the right y-axis (green). Both parameters are presented as percentages, calculated by correction with an uninfected negative control and an untreated positive control. N ≥ 3

**Table 1:** IC50 values for treatment of WT29 & WT94 iMACs and PMMs with miltefosine and amphotericin B. IC50 values were calculated based on the average number of intracellular *L. donovani* or the percentage of infected cells. p-values were calculated using one-way ANOVA followed by Tukey’s multiple comparisons test.

|  | IC<sub>50</sub> | IC<sub>50</sub> |
| --- | --- | --- |
|  | (*L. donovani* per cell) | (% infected cells) |
|  | *Miltefosine* |  |
| **WT29** | 3.85 ± 1.90 uM | 5.83 ± 3.96 uM |
| **WT94** | 3.52 ± 2.28 uM | 4.25 ± 2.36 uM |
| **PMM** | 2.80 ± 0.99 uM | 5.48 ± 3.47 uM |
|  | p = 0.663 | p = 0.813 |
|  | *Amphotericin B* |  |
| **WT29** | 62.1 ± 39.0 nM | 105 ± 77.9 nM |
| **WT94** | 58.0 ± 28.2 nM | 72.9 ± 40.3 nM |
| **PMM** | 12.8 ± 4.76 nM | 19.2 ± 9.35 nM |
|  | p = 0.224 | p = 0.268 |

## Discussion

While great advances have been made in automation, high-throughput screening, and high-content readout in anti-leishmanial drug testing in recent decades (17,35,36), the choice of the appropriate host cell for *Leishmania* remained challenging regarding physiology, practicability, price, and ethics. To date, there is no standard infection model for drug screening for *Leishmania*. Based on our in-house infection assay using PMMs, we developed a new infection model in which human iMACs were infected with *L. donovani* promastigotes. This method is based on the DNA-specific dye DRAQ5 or DAPI, which specifically stains macrophage and parasite nuclei and allows rapid identification and quantification of infected cells. Clear separation based on the size of the parasite, which is several orders of magnitude smaller than the cell nucleus, allows for efficient quantification of infections. Our results show that iMACs are suitable host cells for establishing high infection rates with *L. donovani*.

*Leishmania donovani* is an intracellular parasite that lives and replicates in a phagolysosomal compartment (37). After infection, it induces the formation of parasitophorous vacuoles (PVs), acidic compartments with properties of phagolysosome, in which it develops and replicates. Similar to other models, the intracellular *L. donovani* resides in the phagolysosomal compartment of the iMACs as indicated by LAMP-1 staining (Fig. 4D) (37,38). It has been hypothesised that *L. donovani* arrests the normal maturation process of the phagosome at a stage before the phagosome reaches its full degradative capacity (39,40). Because iPS cells are easy to manipulate on the genetic level our model would allow studying the phagosome maturation after *Leishmania* infection in more detail.

Interestingly, polarisation of the iMACs did not have an effect on the infection rate of the cells. In the natural course of infection, alternatively activated (M2) macrophages are more permissible to *Leishmania* infection than the classically activated and inflammatory M1 macrophages (41,42). As we did not observe an effect on infection rates when infecting polarised cells, we hypothesize *L. donovani* parasites do not have a preference towards of M1 or M2 macrophages for establishing infection, at least *in vitro.* However, this aspect deserves more investigation. CRISPR KO of the surface receptors on iMACs that play a role in the phagocytosis of *Leishmania* parasites such as complement receptor 3, mannose receptors or Fc receptors (43) can provide important information about the entrance of *Leishmania* into the host.

The quantification of infected iMACs to the tested reference drugs correlates with the reference data of infected PMMs from our own laboratory and the scientific literature (33). This is also consistent with what O’Keeffe et al. have previously shown using iMACs as host cells for *L. majo*r*, L. mexicana* and *L. braziliensis* (17). While we achieve consistently high infection rates with this new iMAC-based model, we also report a large variation in the intracellular number of parasites, often reaching up to 50 parasites per host cell. Due to the high infection rates, the number of intracellular parasites per iMAC could be a more accurate parameter for calculating the IC_50_ of anti-leishmanial compounds. Although this model shows promise, the use of high-content imaging techniques as described here is challenging. Inaccuracies in the automated image analysis can either result in an increased false positive signal or not all intracellular parasites are detected. Possibly, the creation of genetically engineered fluorescent parasites could further improve the model. This would allow for easy identification and quantification, as well as live imaging of *Leishmania* in infected iMACs. In addition, fluorescent parasites would be an important step to perform full genome CRISPR screens to find proteins affecting infection. This has so far been limited due to the technical difficulties involved in genetically manipulating macrophages. While THP-1 cells could be used for this, their malignant origin makes them less reliable as physiological models. Fortunately, Navarro-Guerrero found a way to perform CRISPR screening using lentiviral transfection directly in iMACs (44). This approach combined with our infection model could potentially identify important regulators of infection.

As shown, the system allows for intracellular infection and hence future identification of compounds affecting intracellular *Leishmania* replication. Our proof-of-concept experiments with miltefosine and amphotericin B confirmed that targeting intracellular *Leishmania* in iMACs is possible. The next step would be to adapt the system to high-throughput image-based phenotypic screening. The high number of intracellular parasites in our iMAC model opens a great window for performing such screens on a smaller scale, for example in 384-well or even 1536-well plates.

We believe this host cell model provides an alternative that is more ethical than the usage of PMMs and more physiological than THP-1 cells. Human iPS-derived macrophages will allow for more robust anti-leishmanial high-throughput screening given their abundant availability and limited genetic variability, and they will facilitate the study of host-parasite interaction at the genetic level.

## Financial support

This project was funded by the Walter Fischli-Foundation.

## Acknowledgments

We thank Isabelle Fruh, Annick Werner, Monica Cal, Anna Albisetti, Isabelle Claerr and Dominic Trojer for their support in the lab.

## Ethics Statement

Isolation of peritoneal murine macrophages from mice were conducted in accordance with the strict guidelines set out by the Swiss Federal Veterinary Office, under the ethical approval of license number #2374.

## Supplementary Figures

**Fig. S1: *CD marker characterisation of iMAC*** iMACs were harvested from an 84 day-old EB-culture and stained with various CD markers (green) and their isotype controls (grey). Expression is compared to an unstained (black) and an isotype (grey) control. Counts were normalised.

**Fig. S2: *Success of infection vs. EB culture age*** A) iMACs harvested from young (< 1.5 months) or old (>2.5 months) EB-cultures were infected with *L. donovani*, stained with DAPI. Scale bar = 100 µm. B) iMAC infection for iMACs derived from young (dark) vs. old (light) EB-cultures, presented as either number of *Leishmania* per iMAC (grey) or percentage of infected iMACs (green). *P<0.05 was considered significant using unpaired t-test. C) iMACs were harvested from old (130 days) and young (30 days) EB-cultures and stained with various CD markers. Expression is compared between old (blue) and young (orange) cultures and compared to an unstained (black) and an isotype (grey) control. Counts were normalised.

**Fig. S3: *#Leishmania/ iMAC vs. Polarisation*** A) M0, M1 and M2 iMACs were infected with L. donovani, stained with DAPI. Scale bar = 100 um. B) iMAC infection in function of the various subtypes, presented as the number of intracellular parasites per infected iMAC (black) or the percentage of infected iMACs (green). The infection rate was normalised by subtracting the value of the negative control. One-way ANOVA analysis showed no significant difference over polarisation conditions.

## Notes

### Competing Interest Statement

The authors have declared no competing interest.

